# Optimizing contrast in automated 4D-STEM cryo-tomography

**DOI:** 10.1101/2024.02.23.581684

**Authors:** Shahar Seifer, Peter Kirchweger, Karlina Maria Edel, Michael Elbaum

## Abstract

4D-STEM is an emerging approach to electron microscopy. While it has been developed principally for high resolution studies in materials science, the possibility to collect the entire transmitted flux makes it attractive for cryo-microscopy in application to life science and radiation-sensitive materials where dose efficiency is of utmost importance. We present a workflow to acquire tomographic tilt series of 4D-STEM datasets using a segmented diode and an ultra-fast pixelated detector, demonstrating the methods using a specimen of T4 bacteriophage. Full integration with the SerialEM platform conveniently provides all the tools for grid navigation and automation of the data collection. Scripts are provided to convert the raw data to mrc format files, and further to generate a variety of modes representing both scattering and phase contrast, including incoherent and annular bright field, integrated center of mass (iCOM), and parallax decomposition of a simulated integrated differential phase contrast (iDPC). Principal component analysis of virtual annular detectors proves particularly useful, and axial contrast is improved by 3D deconvolution with an optimized point spread function. Contrast optimization enables visualization of irregular features such as DNA strands and thin filaments of the phage tails, which would be lost upon averaging or imposition of an inappropriate symmetry.

## Introduction

Cryogenic Transmission Electron Microscopy (cryo-TEM) traditionally adopts phase contrast imaging as the method of choice for characterization of macromolecular structures at high resolution. Operatively, this involves acquisition of defocused images and computational correction of artifacts introduced by the imaging aberrations that are required for contrast generation. Three-dimensional reconstruction requires a determination of the orientation angles *a posteriori*, typically for tens to hundreds of thousands of molecular poses, in a workflow known as single particle analysis (SPA). Defocus phase contrast imaging was also adopted for tomography, with 3D reconstruction from a series of tilted projections (Medalia et al., 2002). As the view angles are known in advance, tomography enables a direct view of the reconstructed volume, while higher resolution may be achieved by the method of sub-volume averaging (SVA) (Zhang, 2019; Briggs, 2013; Beck & Baumeister, 2016).

In light of the extensive image processing involved, as well as the shortcomings of the weak phase object approximation (WPOA) in describing the electron transmission, it is difficult to interpret local intensities in a quantitative manner by this approach. Structure determination then involves modeling, or in-painting, of a limited number of molecular building blocks, such as amino acids. Another complication is that energy losses on passing the specimen result in chromatic aberration in image formation; each energy loss comes to focus at a different image plane. This is attended in practice by the use of an energy filter in front of the camera, typically to select only the strong “zero-loss” peak and to reject inelastically scattered electrons. Clearly, this imposes a limitation on the specimen thickness that is tied to the mean free path for inelastic scattering (Seifer et al., 2024), which is a few hundred nm in water; this scale is much thinner than most biological cells or even cellular organelles. Accordingly, structural determination of macromolecules at high resolution was originally limited to *in vitro* preparations of purified material.

Scientific interest motivates the observation of functional assemblies *in situ*, however, without removal from the context of the cell or even the organism. This need has led to the development of elaborate protocols for thinning of cryogenically-preserved, vitrified specimens, either using ultramicrotomy (Al-Amoudi et al., 2004; Eltsov et al., 2018) or more commonly focused ion beam (FIB) technology (Villa et al., 2013; Schiøtz et al., 2023). While this facilitates the SVA workflow for in situ structural biology, the three-dimensional cellular context is lost in the thinning process.

Scanning transmission electron microscopy (STEM) offers an alternative modality to wide-field imaging. STEM has been adopted widely in materials science due to the relative ease of image interpretation for strongly scattering samples, but its application to biological imaging is still limited by hardware constraints and lack of familiarity in the community. Early explorations led to its primary application in quantitative mass measurement (Wall & Hainfeld, 1986; Engel, 1978) rather than structure determination. Notably, one such study was performed on hydrated protein filaments under cryogenic conditions (Trachtenberg et al., 1992). More recently, the advantages of STEM were appreciated for tomography of plastic-embedded specimens stained conventionally with heavy metals (Hohmann-Marriott et al., 2009; Wolf et al., 2018). Specifically, STEM is inherently less sensitive than wide-field TEM to chromatic aberration because no real image is formed. Cryo-STEM tomography (CSTET) was introduced for 3D imaging and reconstruction of thicker specimens than could be investigated by energy-filtered phase imaging (Wolf et al., 2014). Quantitative analysis of the scattered electron intensity enables interpretation of the 3D volume in terms of material composition as well as morphology, for example, in distinguishing carbon-rich lipid bodies or heavier inorganic deposits from protein structures (Wolf et al., 2017). A STEM-based SPA reconstruction of the metalloprotein ferritin with limiting concentrations of iron or zinc confirmed that the heavier metal atoms reconstructed with significantly higher intensity (threshold) than the surrounding organic protein (Elad et al., 2017).

Radiation sensitivity is an overriding concern for cryo-microscopy of hydrated organic and biological specimens (Egerton, 2024). Ideally, every scattered or diffracted electron should contribute useful information to image contrast and interpretation. Traditional STEM detectors integrate over a fraction of the diffraction plane: for example, the high-angle annular dark field (HAADF), the bright field (BF), or a variety of intermediates such as annular bright field (ABF) or low-angle annular dark field (LAADF). Azimuthally segmented detectors can be used for differential phase contrast (DPC), by which the realm of high resolution was introduced recently for biological cryo-EM (Lazić et al., 2022). Each of these techniques emphasizes a different contrast mode. At least in principle, they can be acquired in parallel (Seifer et al., 2021), though the possibilities available with commercial hardware are still rather limited. The approach of 4D-STEM utilizes a pixelated detector, i.e., an area camera, to record the entire diffraction pattern for every real space pixel (x,y) as the probe scans (Ophus, 2019). All of the conventional modes may be implemented flexibly by means of “virtual detectors” integrating in software over specific areas of the sensor. Within the bright field region, where there is an overlap of the illuminating field with the scattering, more sophisticated methods sensitive to phase may be implemented. These range from center of mass (COM) analysis, where a lateral shift of the far-field projected illuminating probe is interpreted as a phase gradient acquired on transmission through the specimen, similarly to DPC, to a complete inversion of the projected phase by ptychographic reconstruction. The advent of ultra-fast electron-counting cameras opens the possibilities of ptychographic analysis of biological specimens (Zhou et al., 2020). At higher angles, i.e., in the dark field, 4D-STEM provides a direct map of the differential scattering cross-section integrated along the sample depth (Seifer et al., 2024).

Automated acquisition is a second requirement for effective handling of radiation-sensitive specimens. The various adjustments for mapping, specimen tracking, and focus should overlap the recorded areas as little as possible, and, moreover, the acquisition of large datasets precludes human intervention at the intermediate steps. These considerations are particularly important for tomography, where the same region of interest must be exposed repeatedly as the specimen is tilted. SerialEM (Mastronarde, 2003) is a versatile, open-source software platform that provides all the necessary tools for automated data collection in electron microscopy, including both camera and microscope control, as well as tools for correlation with fluorescence or other imported image maps (Kirchweger, Mullick, Wolf, et al., 2023). Previously, we have integrated a home-built scan generator, SavvyScan (Seifer et al., 2021), with SerialEM for microscope control. The system can record either multiple analog signals from a segmented diode (Opal, El-Mul Technologies, Israel) or conventional photomultiplier-based detectors, or drive a 4D-STEM camera (ARINA, DECTRIS SA, Switzerland). In both cases, the strategy is to return a “diagnostic” image to SerialEM for microscope control while recording the final data to a separate server off-line. The diagnostic image may be simply a single detector output, e.g., from the HAADF detector, or a composite signal such as a sum of diode elements. Using the 4D STEM camera with LiberTEM (Clausen et al., 2023), a number of virtual detectors are also available in real-time. Altogether the system operates similarly to SerialEM in conjunction with frame-based direct detection cameras used in wide-field TEM. By this means, we demonstrate automated 4D-STEM tilt series acquisition taking advantage of the navigation and acquisition schemes built into SerialEM.

## Methods

### Sample preparation

T4-bacteriophages were propagated on E. coli at a multiplicity of infection of 1. After 5h of incubation, the bacterial lysates were removed by differential centrifugation and filtration. Specifically, the bacterial remnants were pelleted for 30 min at 5000g at 4 °C, and the supernatant was filtered using a 0.22 μm pore filter. The T4-phages were pelleted at 35,000 g for 40 min at 4 °C. The T4-phage pellet was then resuspended in 10-100 µL 50 mM Tris pH 8 with 1 mM MgCl2, filtered by 0.22 µm.

The samples were cryo-fixed by plunge freezing using a Leica EM-GP plunger (Leica Microsystems, Vienna). EM grids (Quantifoil R1.2/1.3, with a 2nm continuous carbon layer on 200 mesh copper grids) were glow discharged in a Pelco Easy glow for 60 sec. The blotting chamber was set to 90% humidity and 23 °C. 3 µL T4 phage samples were added to the carbon side of the EM grid, and 2 µL of homemade 15 nm gold fiducials were added to the backside. The grids were either blotted from the back side for 3.5 sec or from both sides for 1.5 sec each, and stored in liquid nitrogen until imaging.

### Acquisition

A Tecnai F20 microscope (TFS) was operated in STEM microprobe mode at 200 kV with 30 µm C2 aperture, for a semi-convergence angle of 0.8 mrad and typically 7-10 pA electron current. The specimen was maintained in a single-axis cryo holder with liquid nitrogen cooling (model 626, Gatan, USA). Images were acquired either by the ARINA pixelated detector with binning to 96×96 pixels in fast mode or the Opal segmented diode detector. Recordings on the pixelated detector were made at a dwell time of 14 µs per probe location in the field of view (operatively 22 µs setting), with 1024124 scan points over a 1.5×1.5 µm field of view, resulting in a dose of 3 electrons/Å^2^. In the case of the segmented detector, the exposure time was set to 5 µs and the current was adjusted for a similar electron fluence. The specimen tilt angles were ordinarily set between -60 and 60⁰ in steps of 2 or 3⁰. The tilt series acquisition was controlled by SerialEM in the dose-symmetric mode, using either a high-angle annular dark field (HAADF / FISCHIONE) detector or any of the segment detectors, whichever is exposed to the beam and provides sufficient contrast. The Record scans were engaged with dynamic focus adjustment during acquisition. Focus correction was engaged only once before the series acquisition. The synchronization of the pixelated detector (ARINA/ DECTRIS) with the scan generator and the HAADF detector was handled with SavvyGate hardware and the SavvyScan software in version 2 (Seifer & Elbaum, 2023). The SavvyScan software also handled the file naming and storage, determined the type of scan, and selection of probe positions within the scans for recording. The graphical user interface of the SavvyScan system sets the acquisition mode and the number of anticipated tilt views, and is operated by clicking the “Start multi” and the “Arm 4D STEM” buttons. A serial number is ordinarily updated with each scan in the tilt series and can be controlled manually to allow repeated scans by user request. The entire collected tilt series data in HDF5 format requires about 50 GB of storage.

### Analysis of 4D-STEM tilt series data

The following steps describe the acquisition protocol and reconstruction of 4D-STEM tilt series. The names of modules in Matlab and Python refer to our shared repository in github (see Data Availability section).

1. Data acquisition: The first step is to acquire HDF5 files that store images of the pixelated detector for every probe location in the scan and for every tilt view. This is handled by SerialEM using our SavvyScan scan engine and digitizer as a client camera. All functions of SerialEM are available, including Low Dose mode and Navigator. In the dose-symmetric tomography mode (Hagen et al., 2017), the tilt views are ordered starting from the center angle and intermittently visiting positive and negative tilt angles. The scan system returns a single image to SerialEM for housekeeping purposes such as focus and tracking. Typically, the HAADF detector is used for this purpose. SerialEM then stores these recorded images in an MRC file. Alternatively, SerialEM can receive and record from an image stream from LiberTEM based on real-time processing from the ARINA camera. In either case, the camera saves the HDF5 data stream to its own server under trigger from the SavvyScan (Seifer & Elbaum, 2023).
2. Optionally: Novena software (DECTRIS) can be used to observe direct results and averaged patterns from the ARINA camera.
3. Optionally: the py4dstem library (Savitzky et al., 2021) may generate images based on 4D-STEM data; one special example is the parallax corrected images (Jupyter notebook hdf5_parallax.ipynb). The tilt series in mrc format may be produced by custom code in Python with libraries of Py4dstem and h5py.
4. Virtual descan: A Matlab function (Arina_trackbeam.m) determines the correlation between systematic shifts in the diffraction disk position and the probe position in the scan. The analysis is performed on the raw data of the pixelated detector, or on a dummy scan over vacuum at the same settings (see explanation below). The function is called by the main processing script (Arinatomo_rings_v9.m) that removes the systemic shifts in the diffraction disk position.
5. Radial and sector segments: The Matlab script (Arinatomo_rings_v9.m) defines 16 segments of annular rings that cover the pixelated detector radii between (1.5+(i-1)*3)±1.5 for i=1..16 (referred as ring1 through ring16). It defines 8 sectors that cover the bright-field (BF) cone (sect1-sect8), and 8 sectors that cover the area outside the BF cone (sect9-sect16). It integrates the electron counts in each of the 32 segments.
6. Center of Mass phase image: The script calculates a COM vector based on the diffraction image in the BF cone area and then calculates the integrated COM (iCOM) image from the vector (Lazić & Bosch, 2017; Seifer et al., 2021).
7. Multiple tilt series: For each of the 32 segments and for the iCOM, the script generates a tilt series of images in the MRC file format, based on the integrated electron count in each probe location at each tilt view. Another script (rings_reorder.m) reorders every tilt series to ascending tilt angles.
8. Tomographic alignment: The scan images are aligned according to a model of rigid body rotation of the specimen. For this purpose, we choose one tilt series that has the clearest contrast (for example, an inner bright field ring or the HAADF series). A number of software packages are available for the purpose, including IMOD (Mastronarde & Held, 2017), AreTomo specifically for fiducial-less alignment (Zheng et al., 2022), and ClusterAlign (Seifer & Elbaum, 2022), which is especially suited for thick specimens. A Matlab script is available (rings_align.m) for cross-correlation alignment and additional corrections similar to the approach in AreTomo. Finally, a script (rings_copyalign.m or rings_clusteralign_translate.m) propagates the same alignment transformations to all other MRC files.
9. Enhanced iDPC: Based on the BF sector counts prepared in step 5 and alignments in step 8, a virtual four-quadrant detector is implemented using a script (sect_iDPC.m) that generates parallax composites of integrated DPC (iDPC), called iDPC1 and iDPC2 (Seifer et al., 2021). iDPC1 is the separate first-order phase contrast part calculated after removing the parallax shifts between the individual quadrant images. iDPC2 is the remaining part of iDPC that ideally concerns defocus-based depth contrast. For samples thinner than the depth of field, iDPC2 or ΔiDPC1 offer variants of iDPC with enhanced contrast (details in the supplementary material).
10. Tomographic reconstruction: We reconstruct all aligned MRC files using ASTRA toolbox libraries, either SIRT-3D (clusteralign_astra_reconstruction.m) or back projection (clusteralign_astra_reconstruction_BP.m) methods (van Aarle et al., 2015; Palenstijn et al., 2011). Instructions are found in (Seifer & Elbaum, 2022). Weighted back-projection was generated using tomo3d (Agulleiro & Fernandez, 2015). Back-projection methods should be followed by deconvolution (Seifer, 2023; Waugh et al., 2020), as discussed further below. The reconstruction results are stored in MRC format.
11. PCA: To generate enhanced contrast, a script (tomos_pca.m) produces a linear combination from the 3D-reconstructions of the virtual ring segments based on principal component analysis (PCA). Typically, the first principal component appears as scattering contrast, similar in texture but inverse to the HAADF, while the second and third principal components appear to emphasize phase gradients. Care should be taken in interpretation of PCA components, however, as it is difficult to associate the physical mechanism of contrast generation with the components of intensity variance in a general way.
12. To generate an orientation map, a script (sect_angle2.m) uses the aligned tilt series of sect9 through sect16 and paints the images according to anisotropy in distribution of the diffraction.

### Virtual descan in 4D-STEM

The Tecnai microscope is not equipped with a hardware descan controller to stabilize the diffraction pattern as the probe moves away from the optic axis. (Descan is in any case an imperfect art.) The problem manifests, for example, as a background shading in DPC or COM images, and is increasingly severe for a larger field of view (lower magnification). For *a posteriori* correction in software, we map the dependence of diffraction disc position (xd,yd) on the probe location (xp,yp) with a low pass filter. Ideally, a linear dependence can be assumed

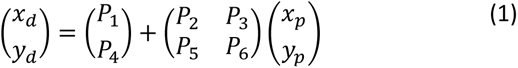

The analysis software extracts the parameters P2,P3,P5,P6 based either on the sampling scan or based on a preliminary scan in vacuum if available. Fig.1 shows the fitted linear model for a scan over a hole in Carbon, and that the alignment does not eliminate the correct beam deflections due to phase gradients.

**Figure 1.**
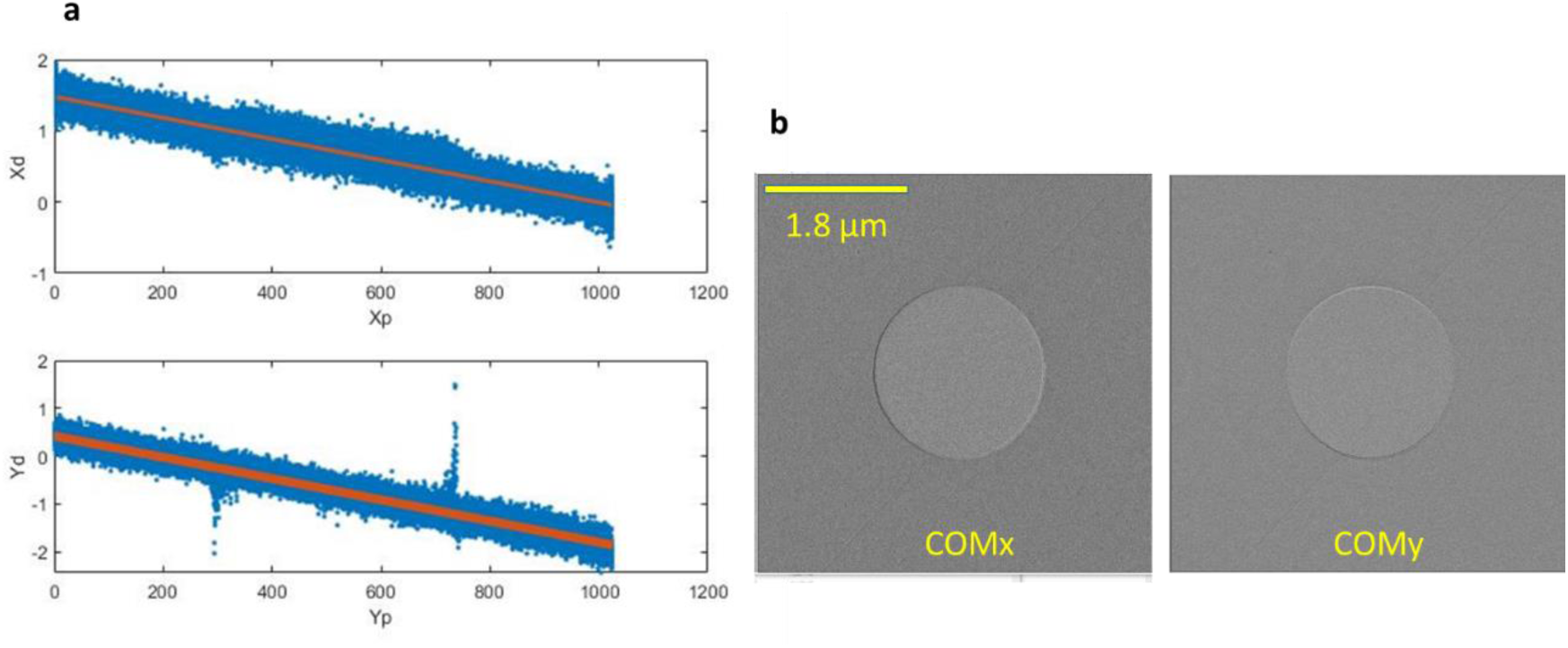
(a) Linear model fitted to the shifts in the diffraction beam position vs. probe position. (b) After the alignment in the beam center based on the linear model, the COM vector shows correct phase gradients at the edge of a hole in carbon.

### Principal component analysis (PCA) in 4D-STEM

Every virtual detector is associate with a different reconstruction file. Together, the annular ring detectors represent a 16-element vector for each voxel. PCA generates the orthogonal eigenvector components from this vector, whose elements are not independent. the first PCA vector typically generates a trivial scattering contrast, while the second and third PCA are sharp in visible contrast. The weighting vectors of PCA1, PCA2, etc., are the vectors obtained in a singular value decomposition of the covariant matrix ordered according to descending singular values. The elements of the covariant matrix are calculated by averaging multiplications of any two elements in the 16-element vector over all the inner layer voxels (correctly recovered) in the 3D reconstructions. The weighting vectors depend on the specimen and optical conditions, but, in general, they take higher contributions from the inner rings of the bright field and the dark field region adjacent to the bright field. Also notable for contrast is the last ring at the edge of the diffraction disk, corresponding to annular bright field (ABF) in a conventional detector configuration.

### Phase images based on either 4D-STEM or quadrant segmented detector

DPC and COM are similar approaches to extract phase information from the specimen based on the deflection of the center of mass of diffracted illumination at phase gradients. These conventional methods are summarized in Fig.2 with reference to the particular hardware configuration that we employed. The respective measurements can be integrated to obtain the proper phase information. We show below that such enhanced DPC analysis is useful in 4D-STEM tomography as well.

**Figure 2.**
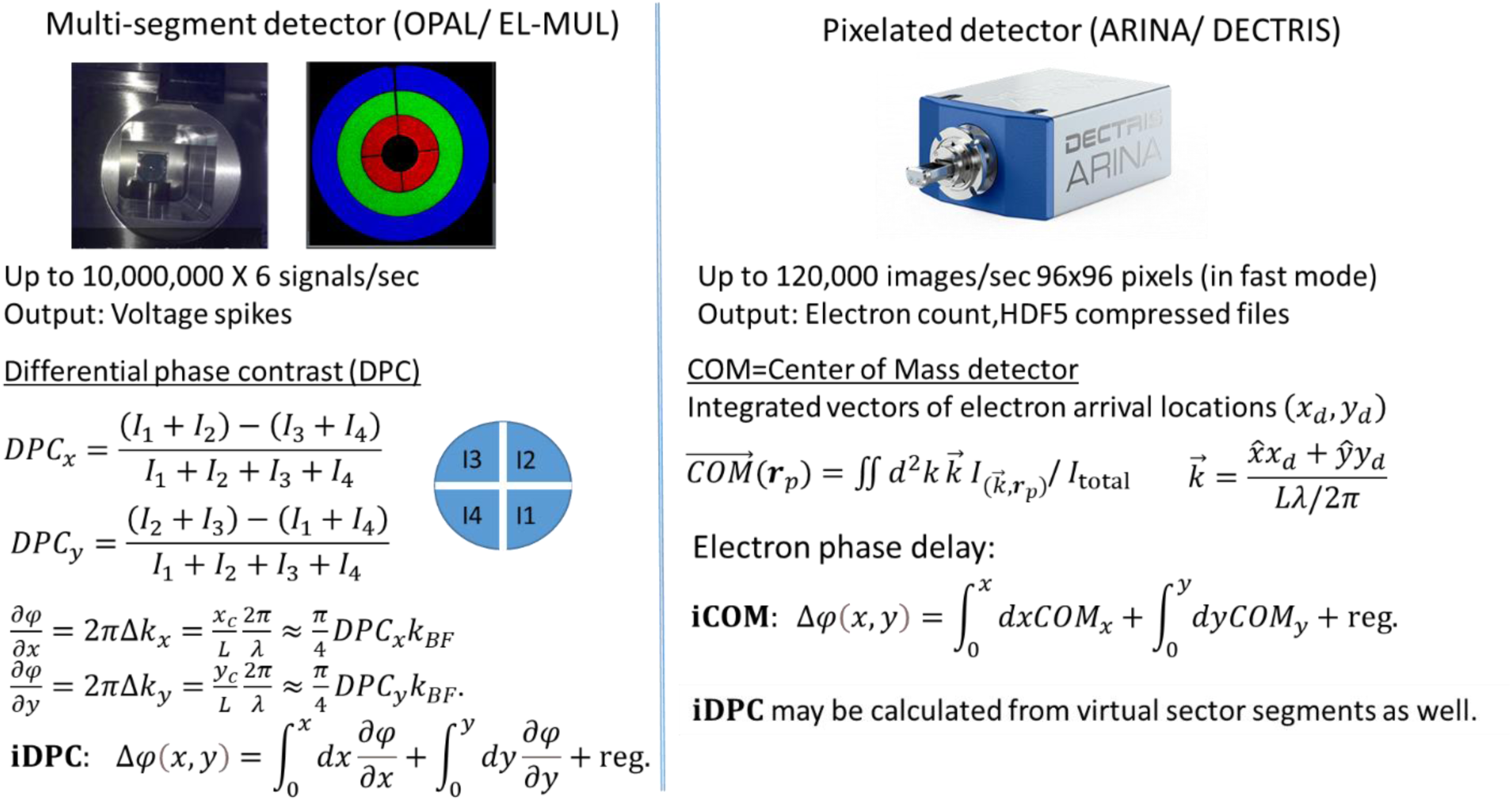
Two options to acquire phase information, using either segmented quadrant detector or pixelated detector. 4D-STEM using the pixelated detector can generate both iCOM and iDPC images.

### Orientation map

Orientation maps display the orientation of lamellar order or preferred diffraction in a 2D plane perpendicular to the incident beam direction. As such, orientation maps are meaningful only in projection scans and not in 3D reconstructions. However, a tilt series of orientation maps is helpful in revealing the order of nano-crystalline structure especially in relation to biological samples. Our orientation maps are generated from the virtual section-detector scans, the files referred as sect9-sect16 in the overall 4D-STEM data processing. An example using a ferritin aggregate may be found in (Seifer & Elbaum, 2023).

### 3D-deconvolution

The idea of 3D-deconvolution has been introduced to STEM as a way to reduce point spread function (PSF) artifacts in weighted-back-projection reconstruction (Waugh et al., 2020). Further automation of the protocol and improvement in speed was described in (Kirchweger, Mullick, Swain, et al., 2023). The method can be justified on the ground of sampling theory using optimal PSF calculation for deconvolution of a back-projection reconstruction (Seifer, 2023).

Suppose that x is the rotation axis of the tilt series and a matrix S* is an operator that generates projections at all tilt angles from a slice of the sample parallel to the yz plane. Back projection is a simple operation of the transpose of matrix S* on the projection data. Optimal reconstruction entails the pseudoinverse matrix operation (SS*)^†^ on the back-projection, which is equivalent to a deconvolution process. The Matlab script (PSF4deconvolution.m) offers a one-time preparation of a PSF file (PSF3x3pipe.mrc) based on the size of the reconstruction and the list of tilt angles. The calculation follows from Eq.25 in (Seifer, 2023), and is reduced to practice as follows

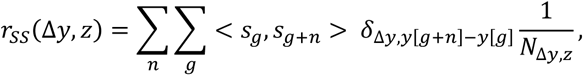

where sn are columns of the matrix S, Δ*y* is the distance from the center of a beam, and z denotes the depth position in the sample. The optimal reconstruction *f̂* is obtained by deconvolution of the back-projection *f_bp_* based on the convolution formula

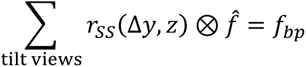

Thus, the 3D PSF is prepared by superposition of *r*_*ss*_(Δ*y*, *z*) at Δ*y* = 0 and ± 1 (3×3×DZ in size) at all tilt views. The remaining steps for 3D-deconvolution are similar to the instructions in (Kirchweger, Mullick, Swain, et al., 2023).

## Results

We utilize the bacteriophage T4 to demonstrate the variety of STEM contrast modes that may be generated using a segmented Opal or pixelated ARINA detectors under conditions suitable for cryo-tomography of fully hydrated, vitrified biological specimens. Diverging from common cryo-EM practice, where many structural insights arise from image averaging, the aim here is to explore the mechanisms for optimal contrast enhancement in single projection images as well as tomographic reconstruction *without averaging*. These possibilities arise from the rich information available in the per-pixel diffraction patterns, which can be exploited flexibly in a variety of different ways.

Fig. 3 presents conventional STEM contrasts using the Opal and HAADF detectors. The HAADF image (contrast inverted) is qualitatively similar to the incoherent bright field image (IBF) generated by a simple sum of the four quadrants. The bright field is subject to parallax image shifts that originate in wave aberrations of the illumination, first and foremost the defocus. The BF image resolution is enhanced by alignment of the images generated by each of the four segments of the Opal detector, essentially “deshifting” prior to taking the sum. The four quadrant segments in the OPAL detector cover diffraction angles between 0.5 and 1.0 mrad, whereas the BF illumination spans up to 0.8 mrad. The small detector areas span a larger depth of field than would a single integrating disk or annulus of similar radius. Image shifts in real space are equivalent to addition of a specific phase to each segment, to compensate the gradients that originate in defocus or other wave apertures. Image shift correction therefore plays a similar role to CTF correction in wide-field TEM, providing a simple implementation of defocus correction. Finally, an iDPC image represents phase contrast.

**Figure 3.**
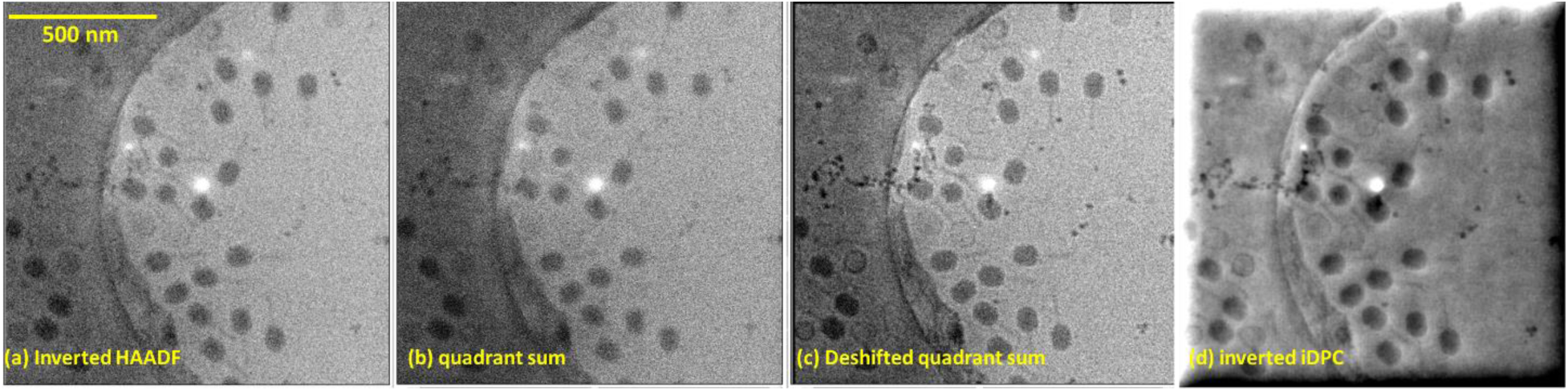
(a) HAADF image with contrast inverted. (b) A simple sum of the OPAL four quadrants that covers part of the BF diffraction disk is slightly less sharp than the HAADF. (c) Sum of deshifted four quadrants is visibly sharper than both HAADF and the simple BF sum. (d) iDPC based on the same four quadrant inputs.

More detailed operations of contrast enhancement and aberration correction are available using the 4D-STEM data from the ARINA pixelated detector. Fig.4 shows projection images of a thin area of the specimen. Virtual detectors are implemented by means of masks on the diffraction plane. Panels (a) and (b) show conventional BF and Annular Dark Field (ADF) images. The gold nanoparticles appear slightly out of focus and their contrast is weak, especially in the ADF, due to Bragg scattering into higher angles that were not acquired. Phages are visible at the perimeter of the hole, though the specimen is dehydrated and empty capsids have collapsed. Parallax correction in 4D-STEM (Fig.4c) imposes image shift corrections at different binning scales (Yu et al., 2021; Savitzky et al., 2021). It may be considered as a more sophisticated analog of the quadrant image shifts demonstrated in Fig 3c, and refocuses the BF image significantly. Contrast is also enhanced by principal component analysis (PCA), which introduces a weighted sum of images acquired by thin virtual annuli detectors covering the diffraction plane (Fig.4d). Unlike traditional bright or dark field STEM, the PCA approach exploits the scattering recorded across the entire camera sensor.

**Figure 4.**
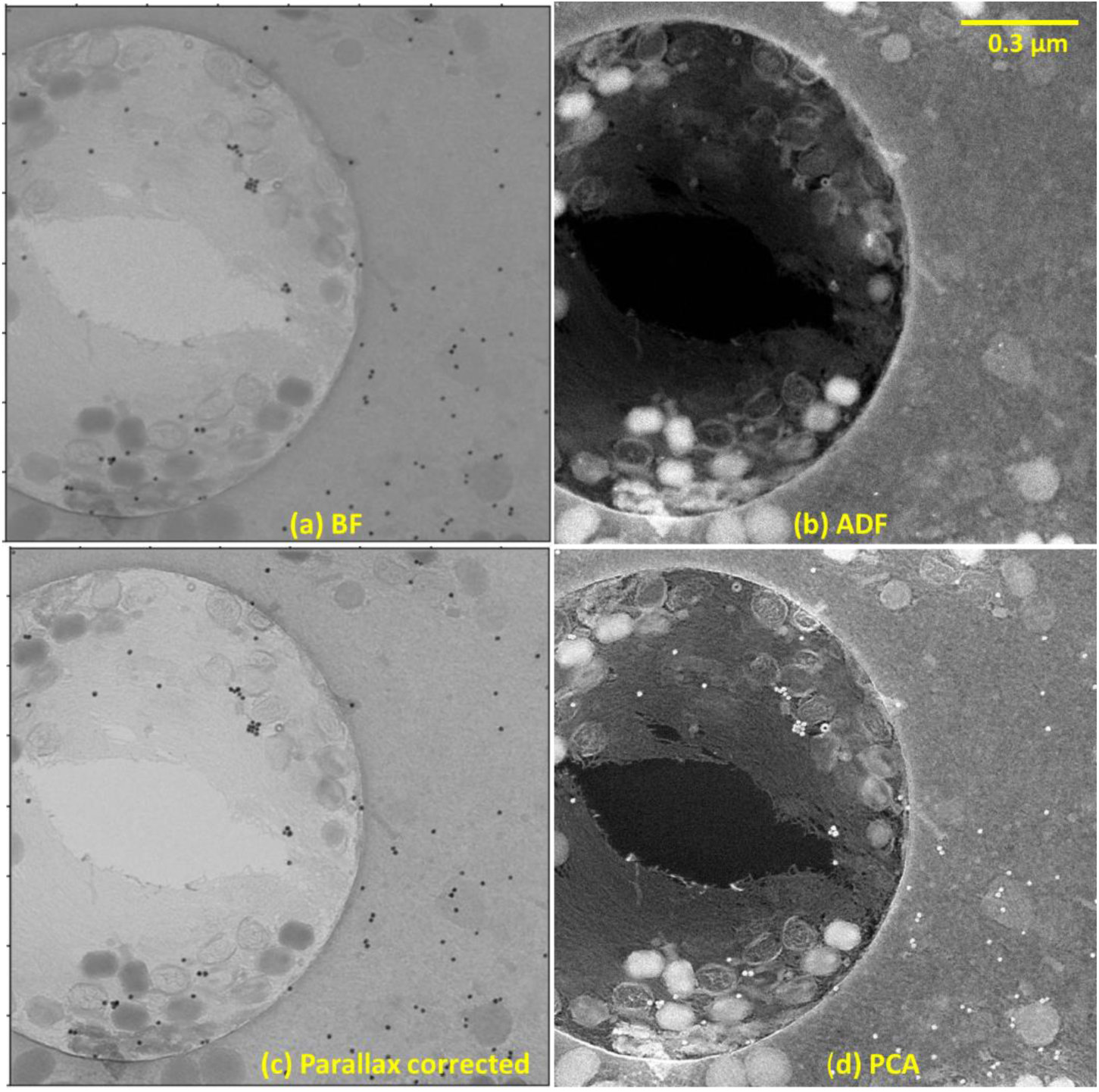
Different contrasts based on (a) BF (b) HAADF (c) Parallax corrected image generated by py4dstem (d) PCA2.

The variety of contrast representations that can be generated in 4D-STEM are also applicable in cryo-tomography. The count of electrons scattered by amorphous biological material may be integrated over the ARINA sensor in concentric thin annuli. Each pixel in real space is then represented by a vector of 16 components rather than a 96×96 pixel array in diffraction space. Each component of the 16-vector serves as a tilt-series that undergoes a 3D reconstruction. Fig. 5 shows the central sections from such reconstructions. The tilt series was acquired between ±44⁰ in steps of 2⁰ for a 200 nm thick sample. A projection camera length was chosen such that the illumination disk fills the inner three virtual annuli. The labels in each panel describe the contrast mechanisms. The top panels present SIRT reconstructions of the three inner rings. Together, these compose the incoherent bright field (IBF) signal, whose image is very similar to that of ring2 alone. The next panels include the iCOM, which integrates a phase gradient signal as described in Fig.2, and virtual annular dark field (ADF), composed of outer segments only.

**Figure 5.**
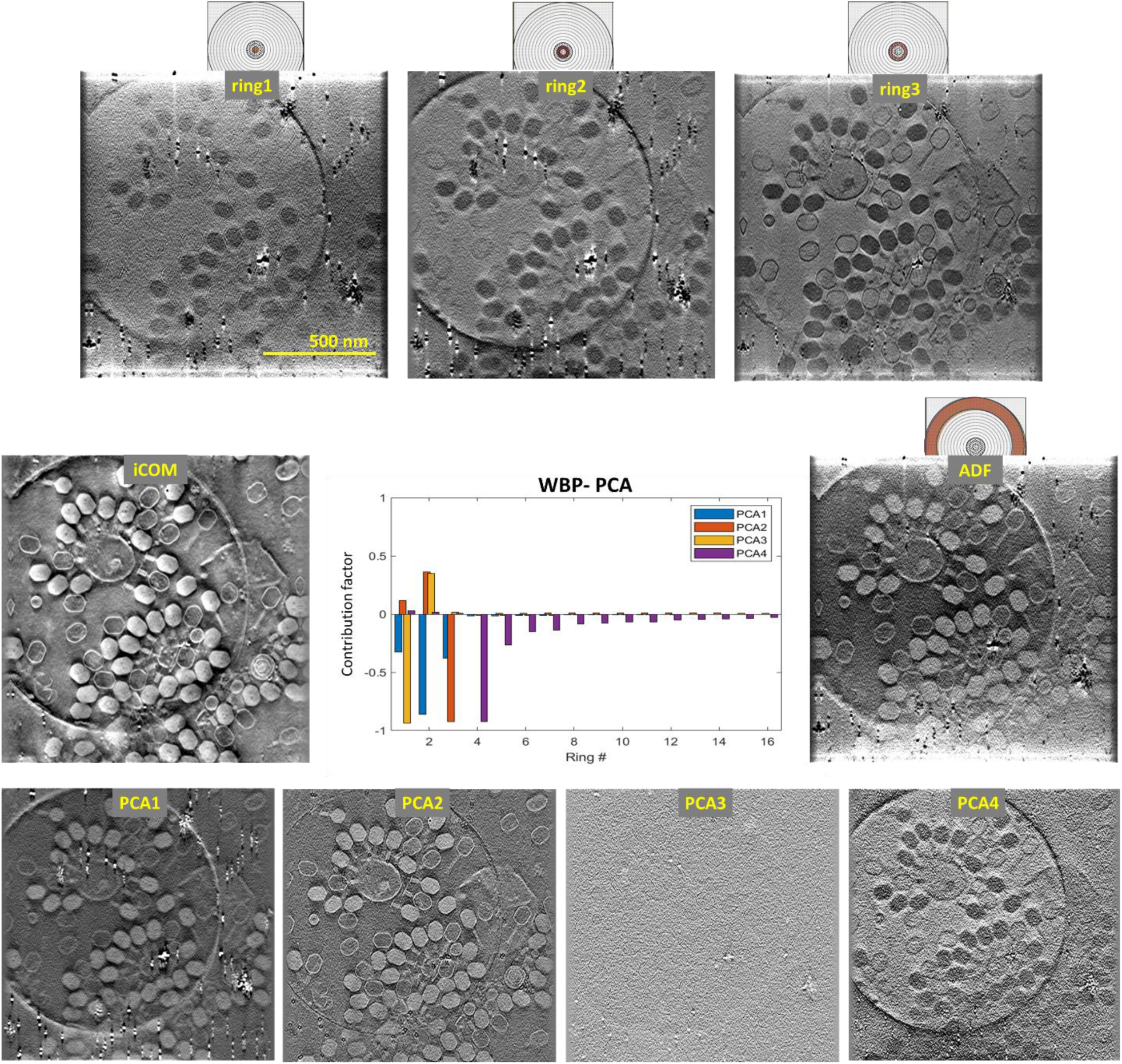
Section of a 3D reconstruction of a 4D-STEM tilt series showing bacteriophages with different contrast rendering from a single dataset of the ARINA pixelated detector. Rings 1-3, iCOM, and ADF are SIRT reconstructions. PCA 1-4 are the superposition of WBP reconstructions from virtual annular detectors with contributions determined as singular vectors from the analysis. The relative contributions to each PCA component appear in the center, normalized such that the sum of squares equals one (vector of unit length).

When examining individual phages, the overall structure of the virions is familiar from both popular and scientific literature (Leiman et al., 2003; Hu et al., 2015). A variety of conformations are visible in all the images. The interiors of the empty capsids show the same contrast as the surrounding aqueous environment. Keeping in mind that the images show a virtual slice through the specimen volume rather than a projection, the image contrast reflects the local scattering potential. The DNA-filled capsids are consistently dark or bright in the bright field or dark field modes, respectively, as are the proteinaceous phage tails. The latter shows two states, extended with a visible base plate and retracted with the central injector exposed. There is also a lipid bilayer vesicle near the center of the support film hole, to which a number of phage are attached; this is a remnant of the purification from lysed bacteria. Phase effects are noticeable in the panel of ring1. According to the theorem of reciprocity, a point-like detector on-axis should produce a STEM image with the same contrast transfer as that of conventional wide-field TEM. Consistently, the phage tails and most of the empty capsid shells disappear. Faint Fresnel fringes appear around some, also consistent with contrast generated by phase interference. The DNA-filled capsids do not entirely disappear, suggesting that their description as weak phase objects is not entirely justified. The phase effects are lost in ring2, since they are averaged over the larger detector area. The ABF image contrast in ring3 is dominated, however, by the phase gradient at material interfaces. The empty capsid shells show very high contrast, and resolution is evidently enhanced. Specifically, striations appear along the tail stalks, and small extensions are seen at the base plate of the extended stalk in the 2D image (as well as others through the reconstructed volume). Also, the gold nanoparticles that had been introduced as fiducial markers for tomographic alignment appear much smaller in the ABF image than in the other modalities. We attribute the strong contrast in the ABF as a signature of small movements of the diffraction disk due to local phase gradients. These cause a change in the coverage of the third virtual ring on the detector, essentially a magnitude of the center of mass (COM) displacements. Notably, the scattering contrast is also represented as the DNA-filled capsids remain dark.

The PCA images in Fig.5 show linear combinations of 3D weighted-back-projection (WBP) reconstructions from the sixteen radial annuli. The contribution factors are determined by the singular eigenvectors in a principal component analysis and displayed in the central panel. The 16 raw components are inter-dependent due to the continuity of the differential scattering cross-sections, as well as phase interference in the bright field. A PCA analysis separates the vectors into orthogonal components and orders them by contribution to the total contrast. The first four PCA components that are shown account for 91% of the total covariance contrast, in which the information is least redundant. The contrast modalities can be recognized directly. PCA1 is the incoherent BF, a positive sum of the three inner rings with most weight to ring2; this represents primarily the scattering contrast. PCA2 is ring3, the ABF, minus a fraction of the signal from ring2. PCA3 is primarily ring1, again minus a fraction of the signal from ring2; this appears as the “reciprocity TEM” BF with the scattering contribution removed. Finally, PCA4 is the composite of the ADF signals; it is visually similar to the scattering contrast of PCA1, but inverted in contrast. Individual long tail fibers can be resolved in the PCA2 image, which provides the sharpest contrast. In general, composites based on the outer ABF ring always represents the phase image with sharp details. PCA assignments are not unique, however. In the case of unweighted back-projection reconstructions from the same dataset (see supplementary material, Fig.S3), PCA3 is the ABF-dominant composition, while PCA1 and PCA2 resemble an incoherent BF contrast. The back-projection PCA span above 99% of the contrast in the first 3 compositions.

Scattered electrons may be collected in sectors within the bright field zone of the detector plane. The virtual iDPC is calculated from a quadrant detector created as a software mask on the ARINA sensor (see Fig.2). Fig.6 compares iCOM and virtual iDPC processed from cryo-4D-STEM images in a tilt series covering a range ±60⁰ with steps of 3⁰. iDPC and iCOM are very similar in detail. Previously (Seifer et al., 2021), we showed that iDPC can be split using parallax decomposition between a phase-related part and a depth contrast, which we denote as iDPC1 and iDPC2 respectively. (Note that iDPC1+iDCP2=iDPC.) Depth of field for beam convergence of 0.8 mrad is approximately 4 μm, which is much thicker than the sample. Yet, the iDPC1 exhibits image corruptions, particularly at low spatial frequencies, that are absent in the iDPC2. iDPC2 is formally related to a linear contrast transfer function (see supplementary material), and we find that iDPC2 is much sharper than iDPC. Close inspection of membranes and empty capsids in iDPC1 reveals a doubling artifact, which coincides with the faint Fresnel fringe at the sharp boundary. Interestingly, the isolated iDPC2 part of the back-projection reconstruction in the XY plane is free of low-frequency distortions. We choose to use unweighted back-projection both as a step in the process toward optimal recovery by 3D deconvolution, and also as a way to stress the character of the parallax decomposition. PCA3 by back-projection (Fig.S4) is particularly sharp in details, and the isolated iDPC2 reconstruction is closer to PCA3 than the iCOM, as indicated by Fourier Shell Correlation. A theoretical explanation for the improvement in clarity is offered in the supplementary material. We also introduce there another phase contrast, ΔiDPC1, which works reliably also without defocus at the object of interest. Essentially, ΔiDPC1 mimics the technique of subtracting two images recorded at different defoci (Donnadieu et al., 2004). iDPC2 is easily implemented after the sector files are generated, and thus it is offered as one benefit of our processing protocol. A second benefit is the ease of acquisition and generation of improved tomograms using 4D-STEM tilt series.

**Figure 6.**
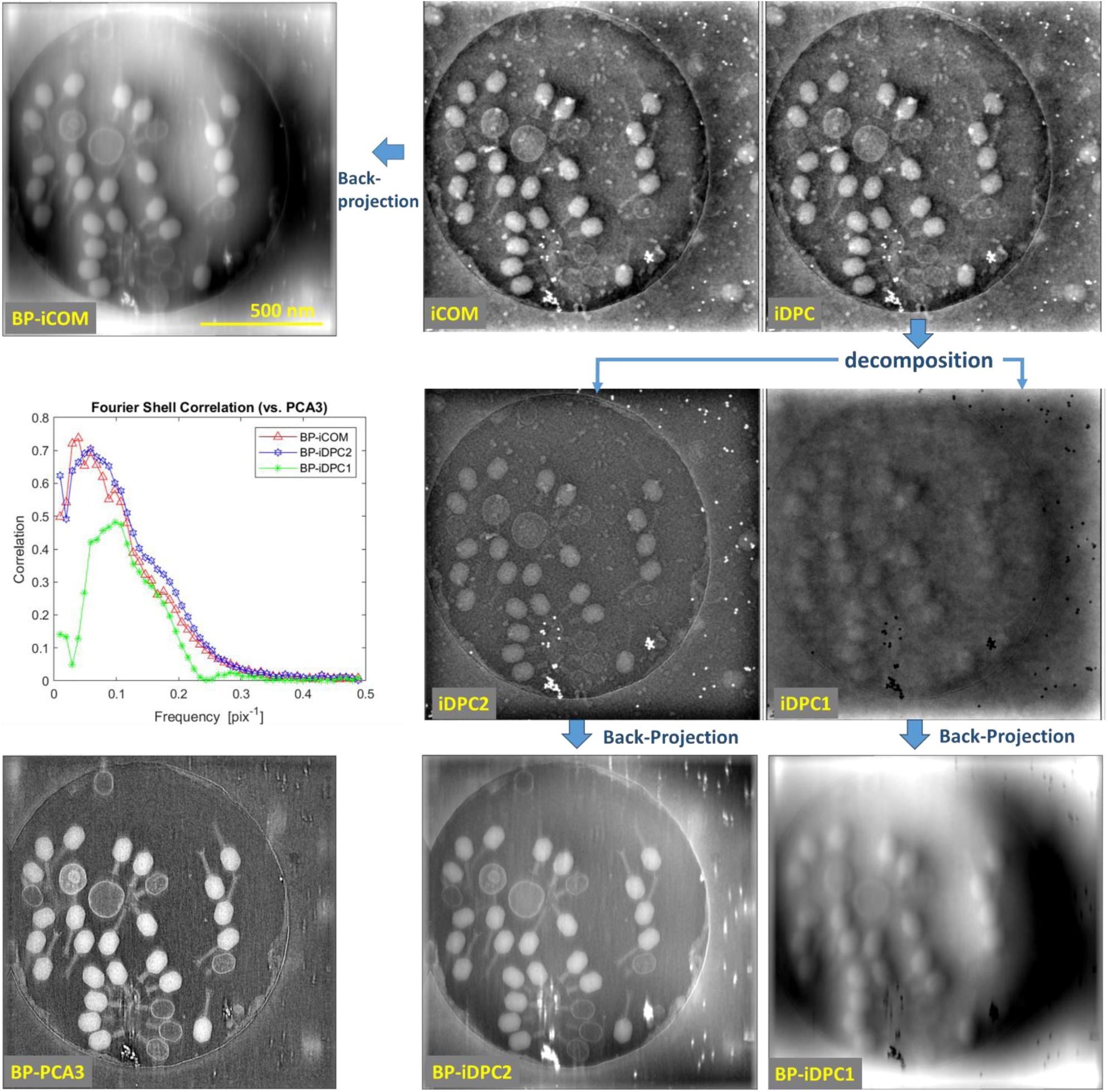
Comparison of iCOM with virtual iDPC in single views; decomposition of iDPC to a parallax-corrected iDPC1 and remaining iDPC2; and back-projection 3D reconstructions. The Fourier Shell Correlation compares the results in 3D to the principal component analysis (BP-PCA3) result as reference. iDPC2 is found most informative, better than iCOM at the high frequencies.

The sophisticated 4D-STEM analyses produce a strong enhancement of contrast in the reconstructed images. However, the axial resolution is still limited by missing wedge artifacts and the small number of projection images in the raw data. We had addressed this previously for conventional STEM by a 3D deconvolution using the synthesized illumination profile as a kernel. As discussed in the method section, 3D deconvolution is essential for optimal recovery after back-projection. Fig.7 shows the results of such processing for the case presented in Fig.6, in comparison to SIRT reconstruction (150 iterations, ASTRA toolbox) and a WBP (SIRT-like 100, IMOD). While a SIRT and WBP reconstructions are much improved in comparison to a simple back-projection, 3D deconvolution after back-projection is still better in terms of sharpness of details. The contrast enhancement is especially significant for the cross-section.

**Figure 7.**
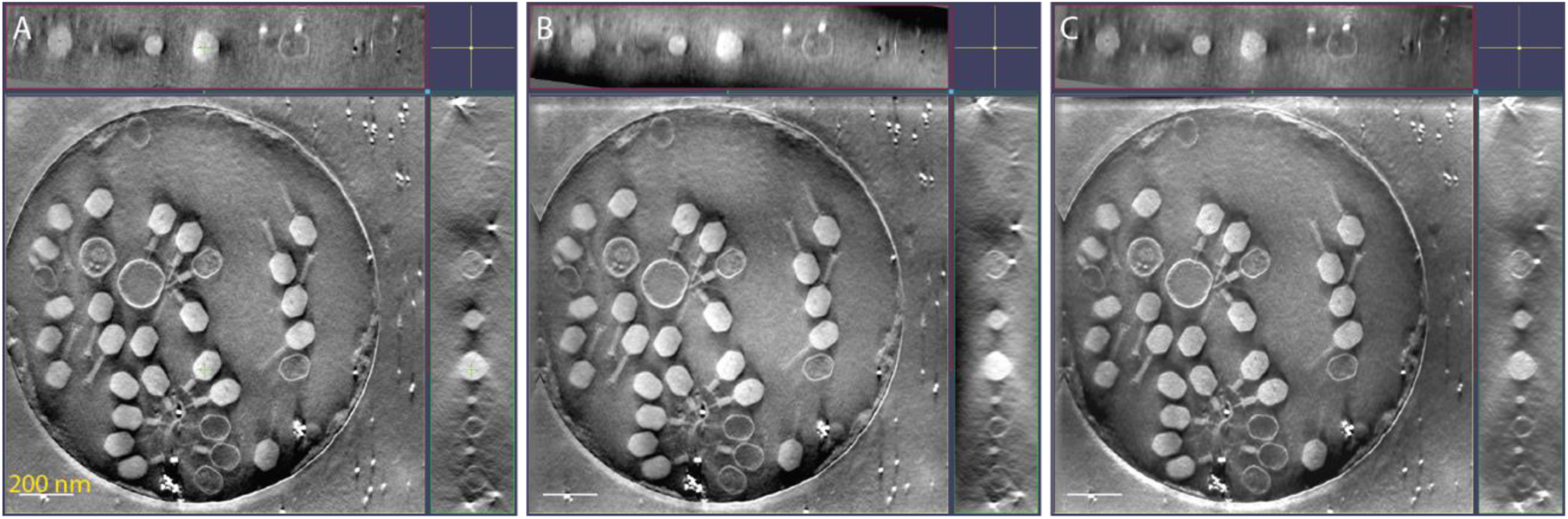
Cross and side section views of 3D reconstruction from iCOM by different methods. (A) SIRT-3D (ASTRA toolbox). (B) WBP with SIRT-like filter (IMOD, 100 cycles). (C) 3D deconvolution (PRIISM) after unweighted back-projection. The green cross in panel A marks the location of the XZ and YZ cross-sections.

This improvement is highlighted when zooming into individual non-averaged virions (Fig. 8 and supplementary material). While iDPC2, iCOM, and PCA analysis are three different contrast mechanisms, PCA2 (cyan) and PCA3 (red) provide different components of the same mechanism. This makes them additive and justifies overlaying them.

**Fig 8.**
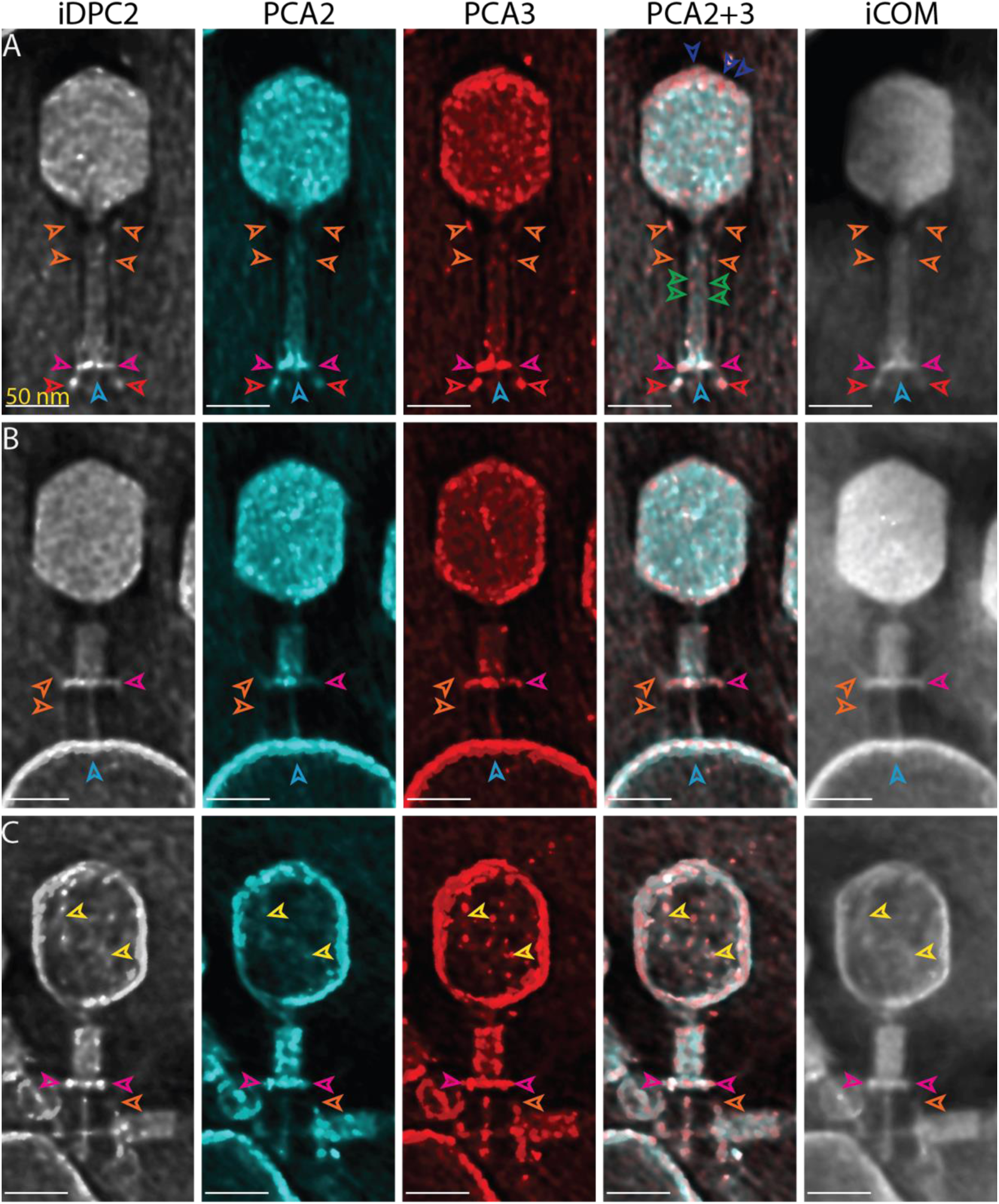
Comparison of iDPC2, PCA2, PCA3, and iCOM after 3D deconvolution of simple back-projection, by zooming in on different confirmations of virions: (A) the full virion, (B) an intermediate virion and (C) an empty virion. The long and short tail fibers, the baseplate, the residual DNA strands, the gp18, the capsid icosahedron, and the gp5 protein cap are annotated as orange, red, magenta, yellow, green, and dark and light blue arrowheads, respectively. 30 slices from the tomograms are shown, covering a depth of 65 nm. The scale bars are 50 nm. Movies of the 3D-volumes and individual slices and additional full virions are found in the supplementary information.

In the full virions (Fig 8A), the DNA-filled head of the virion shows some internal structure, such as fibers and puncta. On the head surface, a punctate structure is observed (dark blue arrowhead), resembling the icosahedron made up of the Hoc, Soc, gp23, and gp24 proteins. The long tail fibers are observed running alongside the tail sheath of the virion (orange arrowheads). The gp18 protein covering the tail (green arrowheads) displays a twisting structure. The baseplate (magenta arrowheads) with the short tail fibers in the stretched-out (red arrowheads) confirmation can be observed. The gp5 protein (blue arrowheads) is visible beneath the baseplate. An intermediate conformation of the virion, which pierces through a membrane vesicle, is observed (Fig 8B). It shows a contracted stalk, which still contains a full head, but a contracted tail sheet. It reveals the inner tube, the end of which is seen inside the vesicle (light blue arrowhead). The long tail fibers are attached to the membrane (orange arrowheads). Empty phages show contracted tail sheets which reveal the inner tube (Fig 8C). Some strands can be observed within the capsid, probably residual DNA (yellow arrowhead). To summarize, while we are focusing on non-averaged examples obtained from a single tomographic tilt series, many molecular details can be observed in the bacteriophage by means of the 4D STEM contrast optimizations, such as protein complex confirmation and DNA fibers. Many of these details would not survive the image averaging based on internal symmetries of the capsid or stalk. Contrast optimization can therefore provide insight on unique features.

## Discussion

Segmented and pixelated detectors promise a new era in cryo-EM imaging. DPC, 4D-STEM, and 2D ptychographic reconstruction have already achieved sub-nanometer resolution by single-particle analysis (Lazić et al., 2022; van Schayck et al., 2023; Küçükoğlu et al., 2024). The combination of ptychographic reconstruction with tomography should be particularly powerful (Pelz et al., 2023). In this work, we demonstrate the potential of 4D-STEM for life science cryo-tomography, leveraging the automation provided by the integration of fast detectors with SerialEM. The data acquisition protocol is compatible with high-throughput approaches and no more difficult than conventional TEM or STEM tomography. Furthermore, we have considered a variety of approaches to extract non-redundant information from the diffraction plane data. This enables the generation of multiple contrast modes, each emphasizing a different physical mechanism of electron scattering or interference, from a single data acquisition. The analyses are relatively simple, based primarily on virtual detectors rather than a complete ptychographic phase retrieval. We also draw insights for a simpler tomography using a quadrant detector, either in hardware or simulated on the camera. DPC may be considered as a first-order approximation to ptychography, and the results are surprisingly impressive. In particular, compensating the defocus aberration by image shifting in real space results in excellent contrast for tomographic reconstruction. The commercially available quadrant diode detectors are still more than an order of magnitude faster than the 4D STEM cameras. Accordingly, we can regard them as four-pixel cameras and call the approach 3.5D STEM.

For both 3.5 and 4D STEM, the essence of phase contrast enters in the difference measurements between half-planes of the diffraction overlapping the illumination. This was predicted long ago (Hawkes, 1978; Rose, 1974). Whereas a phase plate in wide-field TEM generates phase contrast by a π/2 modulation of the unscattered reference wave, similar to Zernike phase contrast, the subtraction operation in DPC STEM is similar to application of a π phase shift on one side. By this argument, the off-axis detection is essential to phase contrast, whether by a DPC or COM strategy. Excellent contrast is observed nonetheless by virtual ABF, as well as by PCA components of the 4D-STEM.

The present data were acquired under conditions of small beam convergence, in which the specimen is much thinner than the nominal depth of field. This is a significant simplification because parallax shifts are approximately uniform throughout the specimen thickness. On the other hand, small illumination convergence limits the lateral resolution to the diffraction-limited probe size, in this case approximately 2 nm. Axial resolution is significantly improved by 3D deconvolution. Features such as the tube inside the phage stalk, individual long-tail fibers, gp5 protein complex, and individual DNA fibers in a near-empty phage head are readily visible without averaging. Especially interesting is the possibility to compare and combine the contribution of different PCA components. With the technical aspects of automated data acquisition in place, future efforts will focus on resolution enhancement and incorporation of ptychographic methods into cryo-tomography.

## Acknowledgements

Bacteriophage T4 were a kind gift of Zeev Barak. We thank Yoav Barak for guidance on the virus incubation and purification. This work was funded in part by the the Weizmann SABRA – Yeda-Sela – WRC Program, and by the European Union (ERC, CryoSTEM, 101055413; Views and opinions expressed are however those of the authors only and do not necessarily reflect those of the European Union or the European Research Council. Neither the European Union nor the granting authority can be held responsible for them.) M.E. is Head of the Irving and Cherna Moskowitz Center for Nano and Bionano Imaging and incumbent of the Sam and Ayala Zacks Professorial Chair in Chemistry. The laboratory of M.E. has benefited from the historical generosity of the Harold Perlman family.

## Data availability

The raw data required to reproduce the above findings are available to download from [https://zenodo.org/records/10679006]. Matlab scripts for data processing are available at https://github.com/Pr4Et/Supplementaries/tree/main/Analysis_4DSTEM_tiltseries

## Notes

### Competing Interest Statement

Decomposition of the iDPC relates to a patent application in which two of the authors are involved (SS & ME). It is referenced in the supplementary material and the text is available at https://patentscope.wipo.int/search/en/detail.jsf?docId=WO2023152734&_cid=P11-LRYVVC-95248-1. There have been no payments or services involved.
The other two authors (PK & KE) have no competing interests.

