## Supplementary Information for "Optimizing contrast in automated 4D-STEM cryo-tomography"

### Contrast approaches

In Fig.S1 we show different contrast approaches based on four off-axis STEM images acquired by the quadrant segments of the OPAL detector for a sample of polymethyl methacrylate (PMMA) sphere on formvar film. The scans were repeated at different defoci as indicated. The image shifts were calculated between all pairs of the 4 images to obtain a characteristic basic shift as indicated at the bottom of the figure. The differential phase contrast (DPC) includes iDPC2, a depth contrast, and iDPC1, a phase contrast. Deshifted sum is an incoherent bright field (IBF) image after image-shift correction of the 4 combined inputs, similar to parallax correction in 4D STEM. HAADF is the input from the high-angle annular dark field detector. ΔiDPC1 is a new phase contrast described below, which mitigates the artifacts in iDPC1.

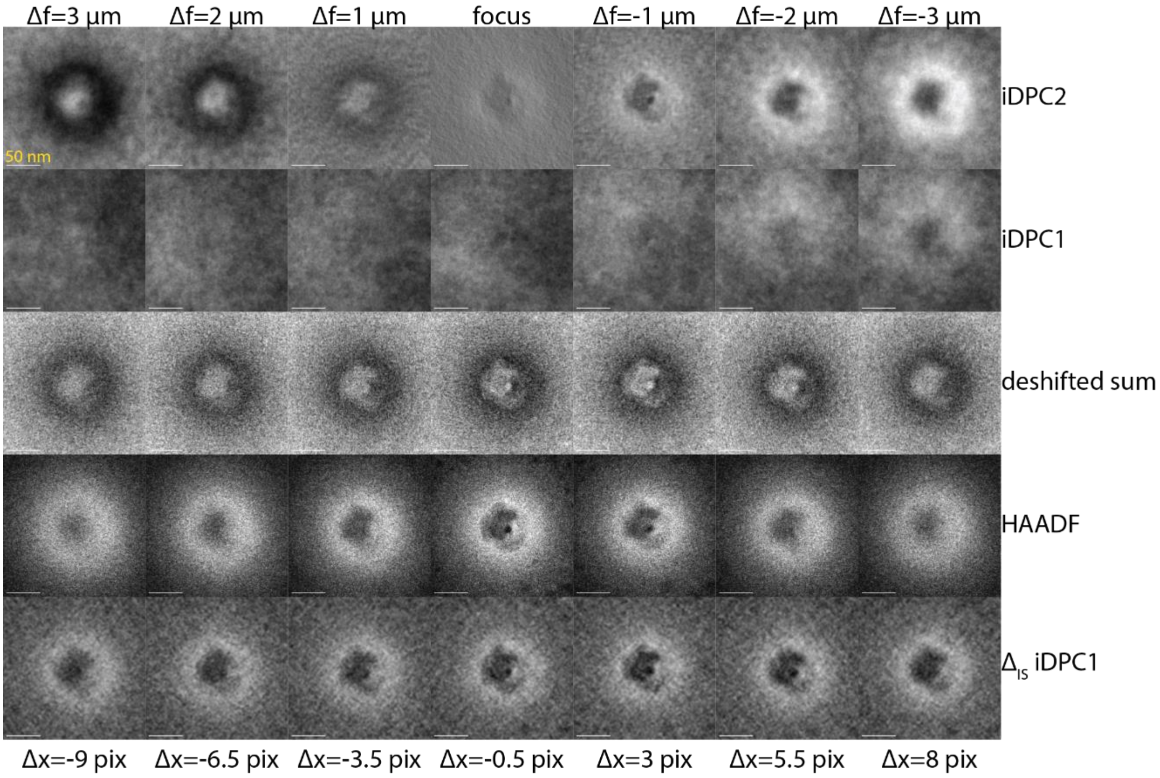


*Figure S1. PMMA sphere in defocus series scanned with OPAL and HAADF detectors, with semi-convergence angle α=4 mrad. iDPC1 and 2 are corrected phase contrast and depth contrast, respectively, based on image shift corrections of the four quadrant detector scans. Image shift corrections compensate low-order optical aberration, i.e., defocus. Spurious phase disturbances are enhanced, however. Differentiation of the iDPC1 result with a slightly modified image shift correction provides a regularized phase image, called $\Delta_{IS}$iDPC1, which is both aberration-corrected and free of disturbances.*

iDPC is known to consist of 3 contributions related to different contrast transfer functions (CTF), which are described in Fourier space as

$$\mathcal{F}\{iDPC\} = CTF_{iS}\,\mathcal{F}\{\varphi\} + CTF_{\varphi^2}\,\mathcal{F}\{\varphi^2\} + CTF_{\varphi^3}\,\mathcal{F}\{\varphi^3\}$$

We decompose iDPC=iDPC1+iDPC2 according to terms with odd and even powers of phase shift φ, so iDPC2 is basically the second-order contribution. In (Seifer et al., 2021) appendix B, we show that $CTF_{\varphi^2} \propto \Delta x$, i.e., the contrast transfer function (CTF) of the second order contribution is proportional to the image shift between the four quadrant images, which basically emerges from features out of focus by the parallax effect. Simulation shows that the influence of defocus on the other CTF is very minor, especially at low convergence angles. Thus, iDPC1 is generated by measuring the image shifts between the quadrant images, removing those shifts by image translations (thus $\Delta x = 0$, and the second order term is eliminated), calculating DPC, and integrating

$$iDPC1 = \int dr \cdot \{DPC(I1_{IS1}, I2_{IS2}, I3_{IS3}, I4_{IS4})\}$$

Ideally, defocus aberration requires the following image shift corrections

$$IS1 = (-\Delta x, \Delta x)$$

$$IS2 = (\Delta x, \Delta x)$$

$$IS3 = (\Delta x, -\Delta x)$$

$$IS4 = (-\Delta x, -\Delta x)$$

Yet, iDPC1 exhibits unwanted disturbances in thick samples, which may be related either to wrap-around of phase in cubic power or to non-conserved properties of the DPC field. A solution to this problem is found by introducing $\Delta_{IS}$iDPC1 calculated as the difference between two nearby image shift correction sets

$$\Delta_{IS}iDPC1 = iDPC1 - \int dr \cdot \{DPC(I1_{IS1+(-1,1)}, I2_{IS2+(1,1)}, I3_{IS3+(1,-1)}, I4_{IS4+(-1,-1)})\}$$

For example, the vector (-1,1) in units of pixel is added to the image shift correction of the first of the quadrant images. The result (see Fig.S1) is a clean phase image with better sharpness, built on the second order phase contribution

$$\mathcal{F}\{\Delta_{IS}iDPC1\} = [CTF_{\varphi^2}\{\Delta x = 0\} - CTF_{\varphi^2}\{\Delta x = 1\}]\,\mathcal{F}\{\varphi^2\} \propto \mathcal{F}\{\varphi^2\}$$

An example of ΔiDPC1 for bacteriophage sample in 4D-STEM is shown in Fig.S2.

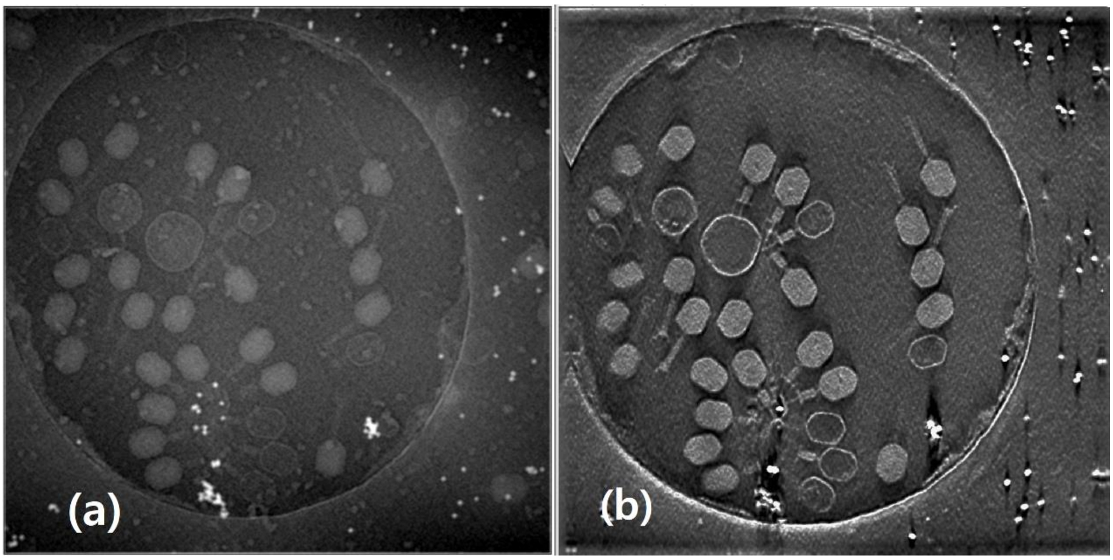


*Figure S2. ΔiDPC1 contrast for bacteriophage sample in 4D-STEM. (a) projection (b) slice of a reconstruction (by back-projection + deconvolution). Compare to Fig 6 for the undifferentiated DPC1.*

We note that parallax corrected differential phase contrast and its derivatives have been found useful also in the light confocal microscope (Elbaum et al., 2023).

### PCA of back-projection reconstructions

Back-projection is a step toward optimal 3D reconstruction. Compositions of back-projection from different rings according to principal component analysis (PCA) are shown in Fig.S3 and Fig.S4. Normally, back-projection (BP) is missing the low frequency suppression provided only in weighted back projection (WBP). However, PCA3 from back-projections is particularly sharp in details.

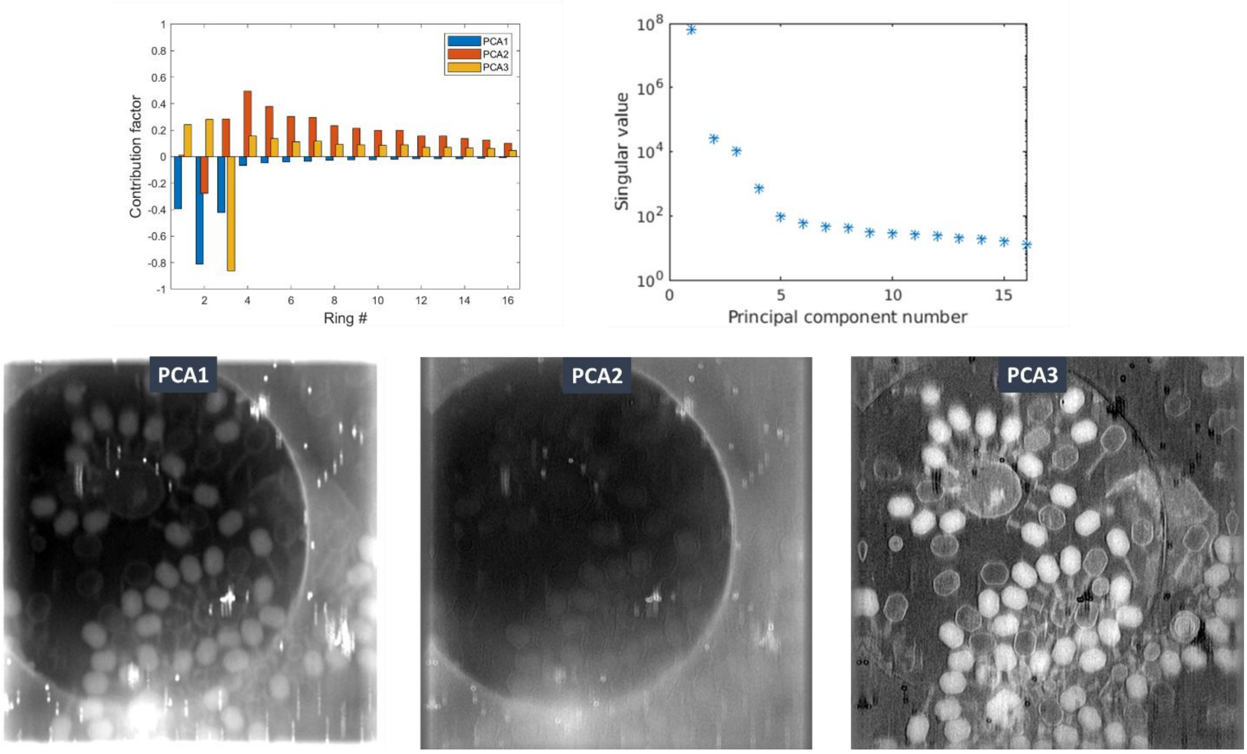


*Fig.S3. Slices of 3D back-projection from virtual annular detectors composed according to principal component analysis (PCA). Example 1.*

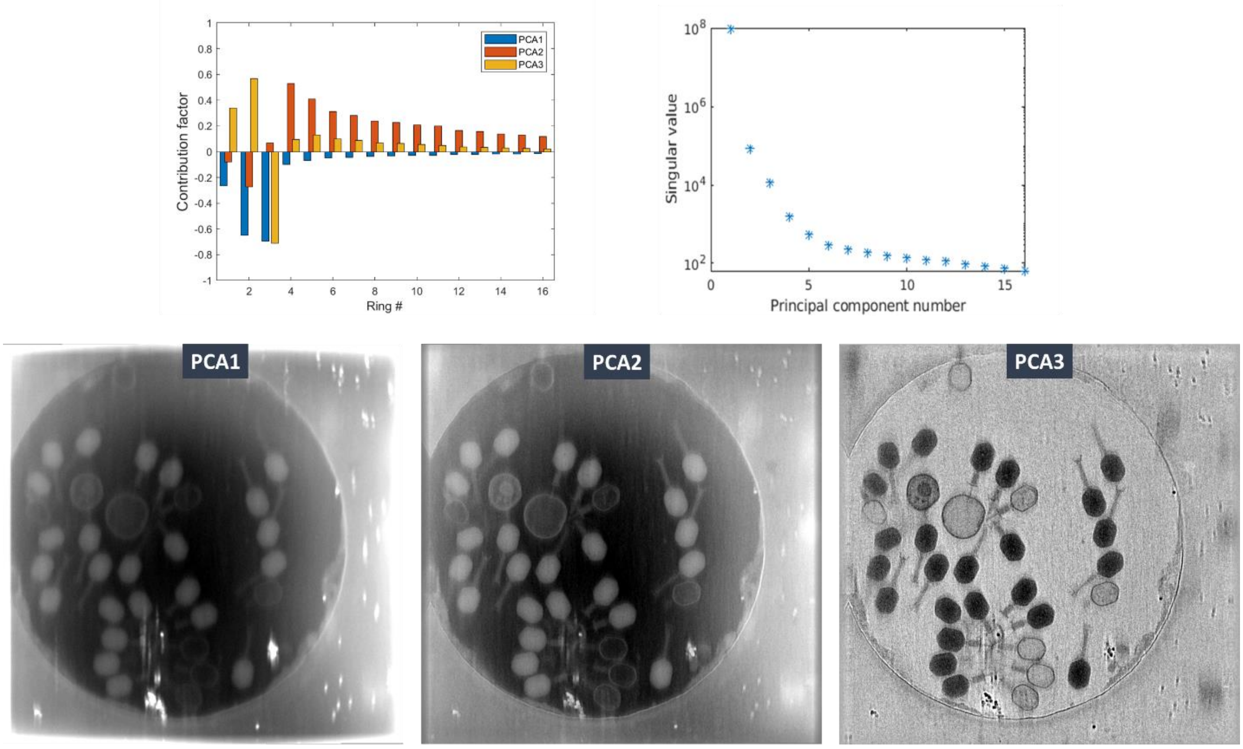


*Fig.S4. Slices of 3D back-projection from virtual annular detectors composed according to principal component analysis (PCA). Example 2.*

### 3D-deconvolution of BP reconstructions in different contrasts

In order to visualize the different contrast modes and the improvements in more detail, we zoomed in on several individual virions and showed 2 slices of the tomograms. We compared three different full virions (Fig S5), a virion in an intermediate stage (FigS6), and a virion in the empty stage (FigS7), and highlighted some macromolecular details.

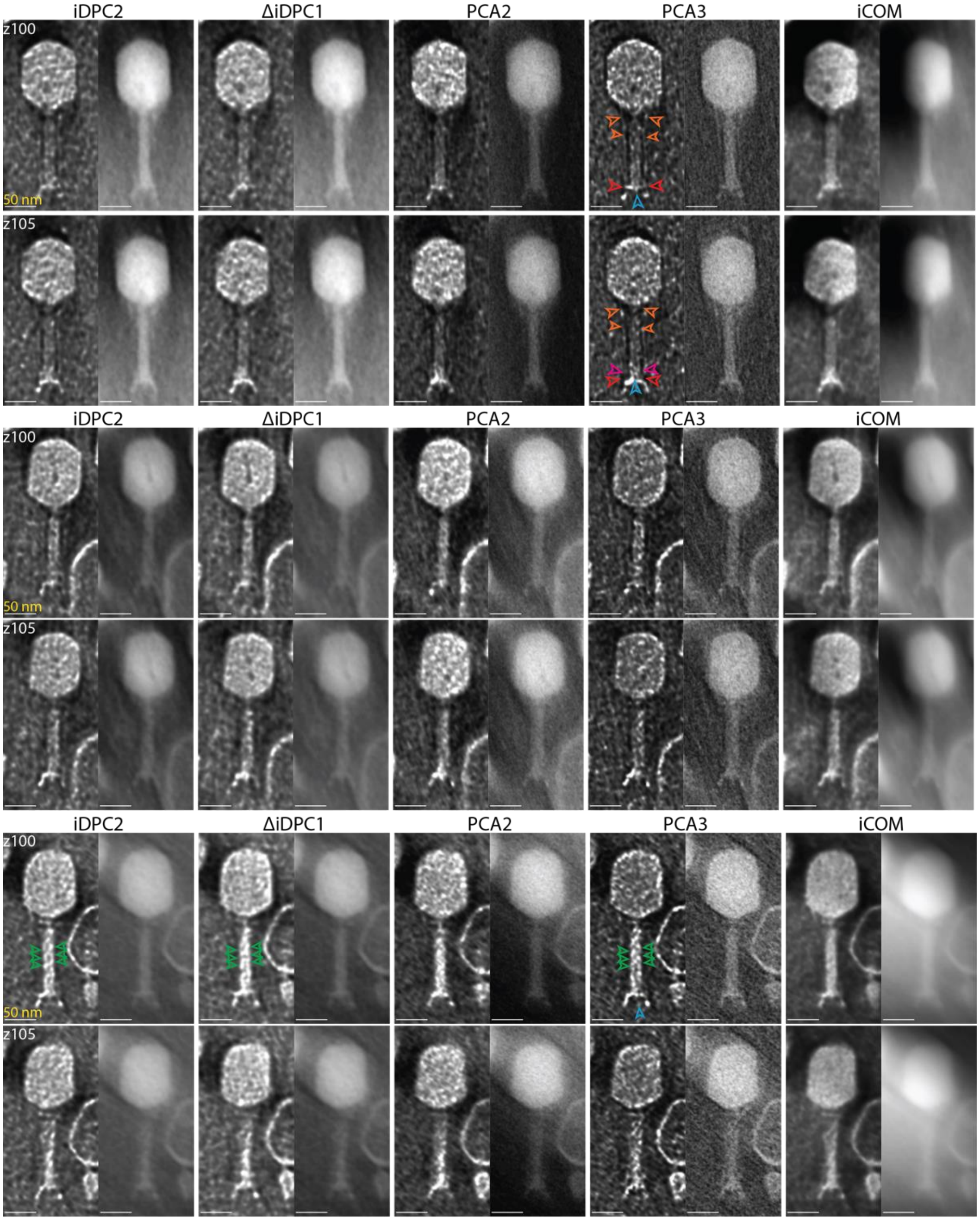


*FigS5: Zoom-in on 3 different virions in the full conformation. Shown are two individual slices per virion. The long and short tail fibers, the baseplate, the gp18, and the gp5 protein cap are annotated as orange, red, magenta, green, and blue arrowheads, respectively. scale bars 50 nm.*

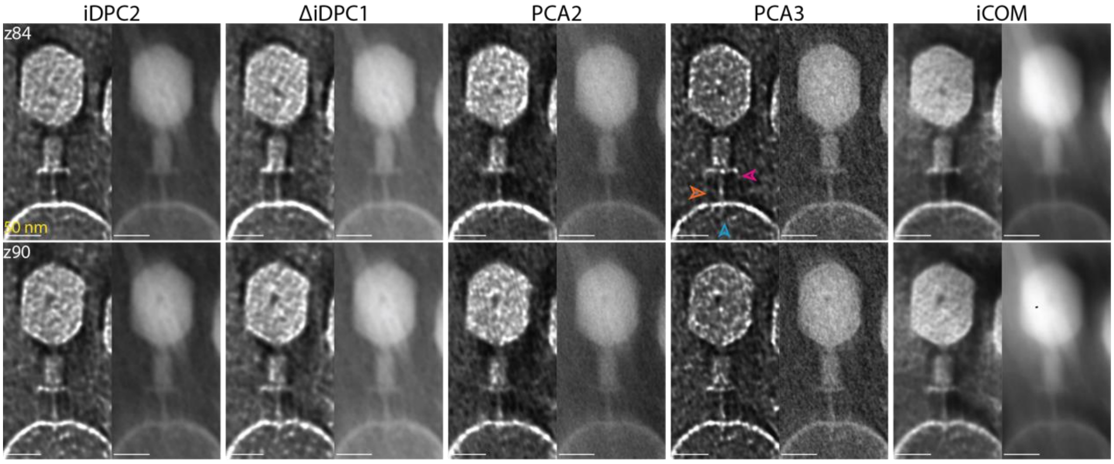


*FigS6: Zoo- in on a virion in an intermediate state piercing through a single membrane vesicle. Shown are two individual slices per virion. The long tail fibers, the baseplate, and the gp5 protein cap are annotated as orange, magenta, and blue arrowheads, respectively. scale bars 50 nm.*

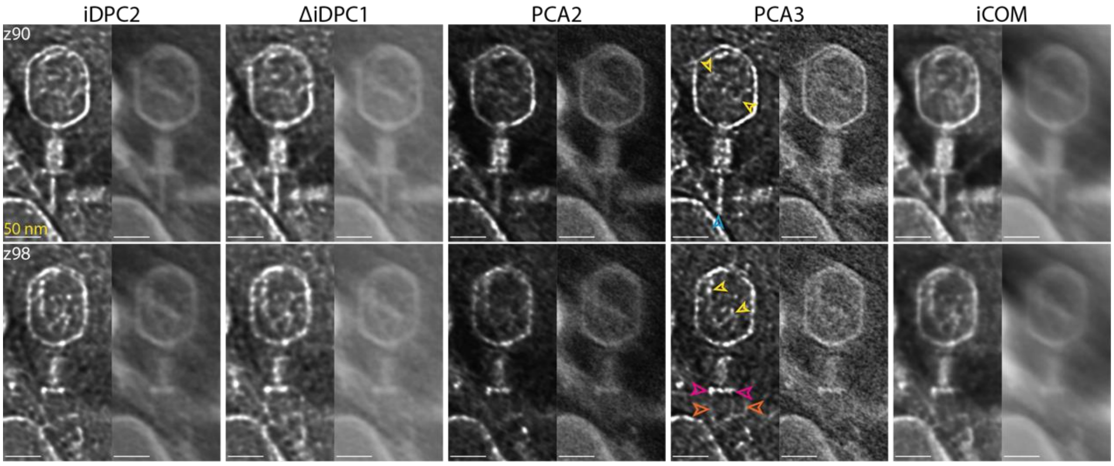


*FigS7: Zoom-in on an empty virion showing residual DNA strands inside the head. Shown are two individual slices per virion. Residual DNA strands, long tail fibers, the baseplate, and the gp5 protein cap are annotated as yellow, orange, magenta, and blue arrowheads, respectively. scale bars 50 nm.*
